# Novel mRNA vaccines encoding Monkeypox virus M1R and A35R protect mice from a lethal virus challenge

**DOI:** 10.1101/2022.11.19.517190

**Authors:** Fujun Hou, Yuntao Zhang, Xiaohu Liu, Yanal Murad, Jiang Xu, Zhibin Yu, Xianwu Hua, Yingying Song, Jun Ding, Hongwei Huang, Ronghua Zhao, William Jia, Xiaoming Yang

## Abstract

The outbreak of Monkeypox virus infection urgently need effective vaccines. However, the vaccines so far approved are all based on whole-virus, which raises safety concerns. MRNA vaccines has demonstrated its high efficacy and safety against SARS-Cov-2 infection. Here, we developed three mRNA vaccines encoding Monkeypox proteins M1R and A35R, including A35R-M1R fusions (VGPox1 and VGPox 2) and a combination of encapsulated full-length mRNAs for A35R and M1R (VGPox 3). All three vaccines induced anti-A35R total IgGs as early as day 7 following a single vaccination. However, only VGPox 1 and 2 produced anti-M1R total IgGs at early dates following vaccination while VGPox 3 did not show significant anti-M1R antibody till day 35. Similar results were also found in neutralizing antibodies and T cell immune response. However, all mRNA vaccine groups completely protected mice from a lethal dose virus challenge and effectively cleared virus in lungs. Collectively, our results indicate that the novel mRNA vaccines coding for a fusion protein of A35R and M1R had a better anti-virus immunity than co-expression of the two individual proteins. The mRNA vaccines are highly effective and can be an alternative to the current whole-virus vaccines to defend Monkeypox virus infection.

## Introduction

Monkeypox virus (MPXV) belongs to the Orthopoxvirus (OPXV) genus of the Poxviridae family, which also includes variola virus (smallpox) and vaccinia virus (VACV) (McCollum and Damon, 2014). Outbreaks of the variola virus had caused millions of deaths in the world till its global eradication in 1980, thanks to the worldwide vaccination with live virus preparations of the infectious vaccinia viruses.

On July 23, 2022, the World Health Organization declared monkeypox outbreak as a global health emergency due to increased cases of Monkeypox infection world-wide (See, 2022). The cease of smallpox vaccination might be one of the reasons causing the current outbreak of MPXV (Reynolds and Damon, 2012) since the two viruses share highly homologous genomes and their antibodies have showed significant cross-protection (Gilchuk et al., 2016; Hooper et al., 2004; Mucker et al., 2022). Currently, there are three pox vaccines available: ACAM2000, a live VACV vaccine (Nalca and Zumbrun, 2010), Modified Vaccinia Ankara (MVA) (Volz and Sutter, 2017) and JYNNEOS (Frey et al., 2014). The JYNNEOS has been recently approved by the U.S. Food and Drug Administration being the main vaccine during this monkeypox outbreak in US.

The vaccinia virus vaccines are replication-deficient in human cells. However, live virus vaccines express many viral proteins. Most of them are still lack of understanding for their functions (Antoine et al., 1998). The attenuated virus vaccines, therefore, remain some safety concerns, especially in immune-deficiency people (Casey et al., 2005; Copeman and Wallace, 1964; Lane et al., 1969).

There are two mainly infectious forms of MPXV: extracellular enveloped virus (EEV) and intracellular mature virus (IMV). Subunit vaccines that use selected recombinant viral proteins from EEV and IMV may have better safety than the live virus vaccines. Vaccination with the E. coli-expressed A27L, a truncated IMV surface protein, protected mice from a lethal challenge of VACV (Lai et al., 1991). Vaccination with recombinant viral EEV proteins B5R or A33R also protected mice from lethal challenge of smallpox (Galmiche et al., 1999). Hooper et al. found that mice vaccination with DNA encoding L1R (an IMV protein) and A33R induced neutralizing antibodies against L1R and IgGs against A33R. Combination of the above two genes was more effective than either gene alone in protecting mice against lethal challenge of VACV (Hooper et al., 2000). They also found vaccination with DNA encoding vaccinia virus genes (L1R, A27L, A33R, and B5R) protected rhesus macaques from severe disease post the lethal challenge of MPXV (Hooper et al., 2004). However, Kaufman et al. found adenovirus-expressing L1R effectively protected mice from a lethal systematic infection and behaved better than the combination of L1R, A27L, A33R, and B5R while both L1R and A33R are required against intranasal infection (Kaufman et al., 2008). Therefore, selection of viral antigens and combination strategies for better vaccine protection against the MPXV remain to be explored.

Since the overwhelming success of mRNA vaccine in defeating SARS-Cov-2 (Baden et al., 2021; Polack et al., 2020), mRNA vaccines with lipid nanoparticle delivery system have raised great interest. The benefits of mRNA include it is fast to produce in cell-free system, no risk of host genomic integration and can stimulate both humoral and cellular immune response (Pardi et al., 2018). In the current study, we have developed mRNA vaccines expressing MPXV EEV protein A35R (homolog of A33R in VACV) and an IMV protein M1R (homolog of L1R in VACV). Those vaccines were tested for their humoral and cellular anti-VACV immunity as well as their protection against the lethal dose of viral infection in mice. We found that novel mRNA vaccines expressing a fusion protein composed of a truncated form of A35R and a full length M1R can provide strong immunity and protection against the poxvirus.

## Materials and Methods

### Cell lines and virus

Vero cells and 293T cells were originally obtained from American Type Culture Collection and cultured in routine complete dulbecco’s modified eagle medium with 10% FBS and 1% penicillin & streptomycin. Vaccinia virus Western Reserve (WR, VR-1354) was obtained from ATCC and propagated in Vero cells. The infected cell lysates and supernatants were collected and ultracentrifuged followed by virus titers quantification using routine plaque assay.

### Antigen sequences and plasmids

M1R and A35R protein sequences from MPXV strain Zaire79 were used to reversely translate to their coding sequences by GenSmart™ Codon Optimization (GenScript). The A35R extracellular domain was fused with M1R by a peptide linker was also synthesized. The corresponding DNA sequences were synthesized by Azenda Life Sciences and cloned into pUC vectors with an upstream T7 RNA polymerase promoter and downstream polyA sequences. The correction of sequences was confirmed by DNA sequencing.

### In vitro transcription

Plasmids were linearized by enzyme BSPQI (Vazyme, Nanjing) as template for mRNA preparation. Briefly, reactions containing T7 RNA polymerase, NTPs, DNA template and RNase inhibitor were incubated at 37 degree for 2 hours followed by mRNA purification using a commercial kit. The integrity of mRNA was verified by agarose gel analysis.

### Transfection and western blot

MRNA transfection was followed by protocol of Lipo3000 with little modification. Briefly, 293T cells were plated in 24-well plates in OptiMEM and transfected with 800 ng mRNA/ well. At 16 hours post transfection, cells were scraped in SDS-loading buffer (Beyotime, Shanghai) and boiled at 95 degree for 5 minutes. Antibodies used in this study were below: M1R Human Mab (OkayBio, R403k5), 1:1000; A35R Mouse Mab (Sino Biological, 40886-M0017), 1:1000; GAPDH Mouse Mab (Sangon, D190090), 1:2000. HRP-goat anti human IgG (Sangon, D110150), 1:5000; HRP-goat anti mouse IgG (Sangon, D110087), 1:5000.

### LNP-encapsulation

MRNA was encapsulated within a lipid formulation (Chen et al., 2012; Leung et al., 2012) to form mRNA-LNPs using a Nanoassemblr^®^ microfluidic device (Precision Nanosystems Inc., Vancouver, Canada). The residual Citrate salt and Ethanol were removed from raw mRNA-LNP after dialysis. Finally, the dialyzed mRNA-LNP solution was preserved at −80°C with a cryoprotectant.

### Mice vaccination

Balb/c mice (female, 7-8 weeks old) were obtained from Vitalriver (Beijing). Ten micrograms LNP-mRNA in 100 μl was intramuscularly injected per mouse and the mice were boosted at 14 days post 1^st^ vaccination.

### ELISA

A35R protein (OkayBio, C1620) and M1R protein (OkayBio, C1624) were diluted to 5 μg/ ml by ELISA coating buffer (Elabscience, E-ELIR-003) and added into 96-well plates for overnight coating at 4 degree. Washing buffer was used to wash out any uncoated proteins before incubation with blocking buffer at room temperature for 1 hour. Blood, collected from different timepoints, were centrifuged and serum were added into wells with different dilution. After incubation, the sera were washed out and HRP-goat anti-mouse IgG (Sangon, D110087) was added for final optical density analysis in a MD microplate reader.

### PRNT

The plaque reduction neutralization test (PRNT) was used to examine neutralizing antibodies. Briefly, serum was heat-inactivated at 56 degree for 30 minutes and then 100 μl serum with different dilution was incubated with 200 PFU virus (VACV-WR) in 100 μl in 96-well plates at 37 degree for 1 hour. The above virus-serum mixture was then added into Vero cells at 37 degree for 1 hour and the plates were shaked every 15 minutes to assure virus fully accessing to cells. Five hundred microliters of 1% methycellulose was added to each well and cells were cultured for 2 days for plaque formation. Cells were fixed followed by stained with 1% crystal violet.

### Flow cytometric analysis

Spleens from mice at day 30 post 1^st^ vaccination were collected and grinded with filters. Cells were suspended in RPMI1640 (2%FBS) medium followed by red blood cells removal. Cells were plated in 96-well plates with 1×106 cell/ well and cultured for 18 hours with treatment of proteins M1R or A35R. Cells were centrifuged to be resuspended in medium containing GolgiStop. After 5 hours stimulation, cells were centrifuged and washed with PBS followed by staining in a PBS buffer containing cell surface markers at 4 degree for 30 minutes. Cells were then centrifuged, washed and fixed in fixing buffer from an intracellular staining kit (BD Bioscience, 554714) at 4 degree for 1 hour. Cells were then centrifuged, washed twice and incubated with intracellular cytokine staining reagent. Cells were then centrifuged and washed followed by flow cytometric analysis.

### Virus challenge

At day 36 post vaccination, mice were intranasally inoculated with 1×10^6^ PFU VACV-WR virus/mouse. Body weight and clinical symptoms were measured every day and mice were sacrificed until weight loss more than 15% or at 9 days post challenge. Lungs were collected and preserved in complete DMEM.

### Viral load in lungs

Lungs were grinded in a tissue homogenizer followed by 3 times freezing-melting to release virus from cells. After centrifugation, supernatants with different dilutions were added to Vero cells for plaque assay.

### Statistical analysis

Data were performed by Graph pad software and One-Way ANOVA was used to analysis differences where “*” indicating p < 0.05, “**” indicating p < 0.01, “***” indicating p < 0.001, and “****” indicating p < 0.0001.

## Results

### MRNA design and in vitro protein expression

As shown in Fig. 1A, totally four codon-optimized mRNA were synthesized, which encode MPXV full length A35R, M1R, integral extracellular domain of A35R fused with full length of M1R (SP-A35R IECD-M1R) and a shorter extracellular domain of A35R fused with full length of M1R (SP-A35R sECD-M1R), respectively (Fig.1A). Protein expression of the 4 mRNAs was confirmed by mass equally mRNA transfection into 293T cells followed by respective incubation with anti-M1R and anti-A35R antibodies (Fig. 1B). All proteins were expressed and detected by western blotting. The mRNA coding for SP-A35R sECD-M1R expressed higher protein level than SP-A35R IECD-M1R in 293T cells.

**Fig. 1.**
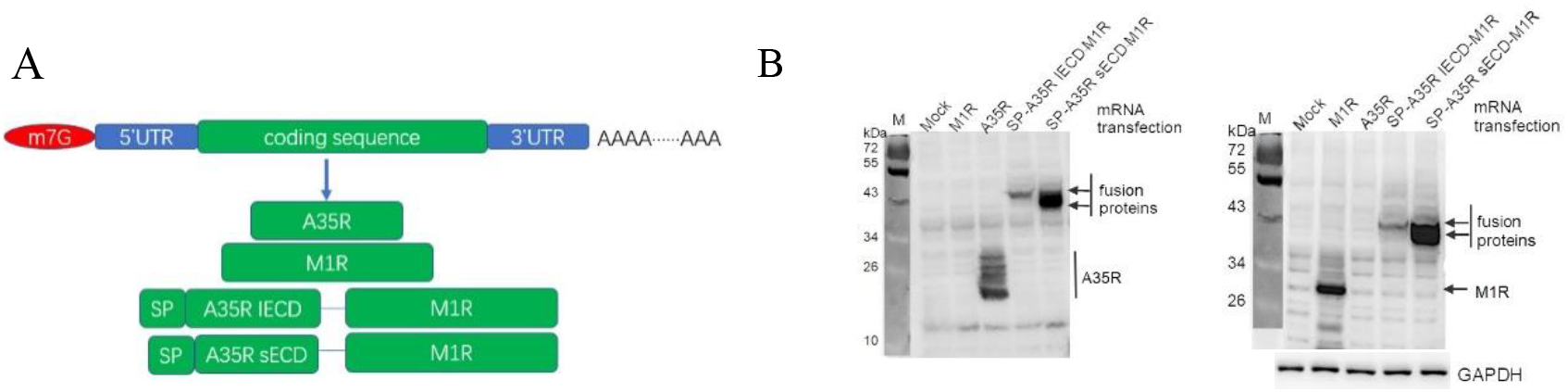
mRNA design and protein expression in transfected 293T cells. (A) mRNA design. MPXV A35R and M1R sequences were codon-optimized and inserted into the region between 5’UTR and 3’UTR. Two mRNAs encoding fusion proteins composed of an A35R extracellular domain and a full length of M1R. SP: signal peptide; IECD: integral extracellular domain; sECD: an extracellular domain lacking stalk region. (B) protein expression measured by western blotting. mRNA was mass-equally transfected into 293T cells and at 16 hours post transfection cell lysates were loaded into SDS-PAGE gels for western blot analysis. left panel: anti-A35R antibody incubation; right panel: anti-M1R antibody incubation.

### A35R and M1R specific total IgGs and neutralizing antibodies

Based on the above results, we developed 3 mRNA-LNP complexes. As shown in Figure 2A, VGPox 1 and VGPox 2 are LNP encapsulated SP-A35R IECD-M1R and SP-A35R sECD-M1R, respectively. VGPox 3 contains a combination of two individual LNP-mRNA complexes coding for A35R and M1R, respectively. The above LNP-mRNA complex were injected intramuscularly into mice in 4 groups (Fig. 2A). Fig. 2B showed mice were vaccinated at day 0 and day 14. The blood was sampled at day 7, 13, and 35 for antibody analysis. In separate groups, the spleens were sampled at day 30 following vaccination for cellular immunity analysis. At day 36 post-vaccination, mice were intranasally inoculated with a lethal dose of VACV-WR (1×10^6^ PFU/ mouse) (Smee et al., 2001; Wyatt et al., 2004). Anti-A35R antibodies were induced as early as day 7 post vaccination in all 3 vaccine groups and increased at day 13 and 35 (Fig. 2C). In contrast, anti-M1R antibodies were only found in VGPox 1 and VGPox 2 not VGPox 3 groups at day 7 and 13. The M1R IgG was only detectable at Day 35 in low dilution sera from VGPox 3 group (Fig. 2D). As shown in Fig. 2E left panel, sera from VGPox 1 and VGPox 2 but not VGPox 3 groups presented neutralizing activities against live virus (VACV-WR) at day 13 and VGPox 2 behaved better than VGPox 1 at 1:100 dilution. The sera from all 3 mRNA vaccines can partly or nearly fully neutralize virus at 1:50 dilution at day 35 while VGPox 1 and VGPox 2 both showed significantly higher neutralizing ability than VGPox 3 at 1: 500 serum dilution.

**Fig. 2.**
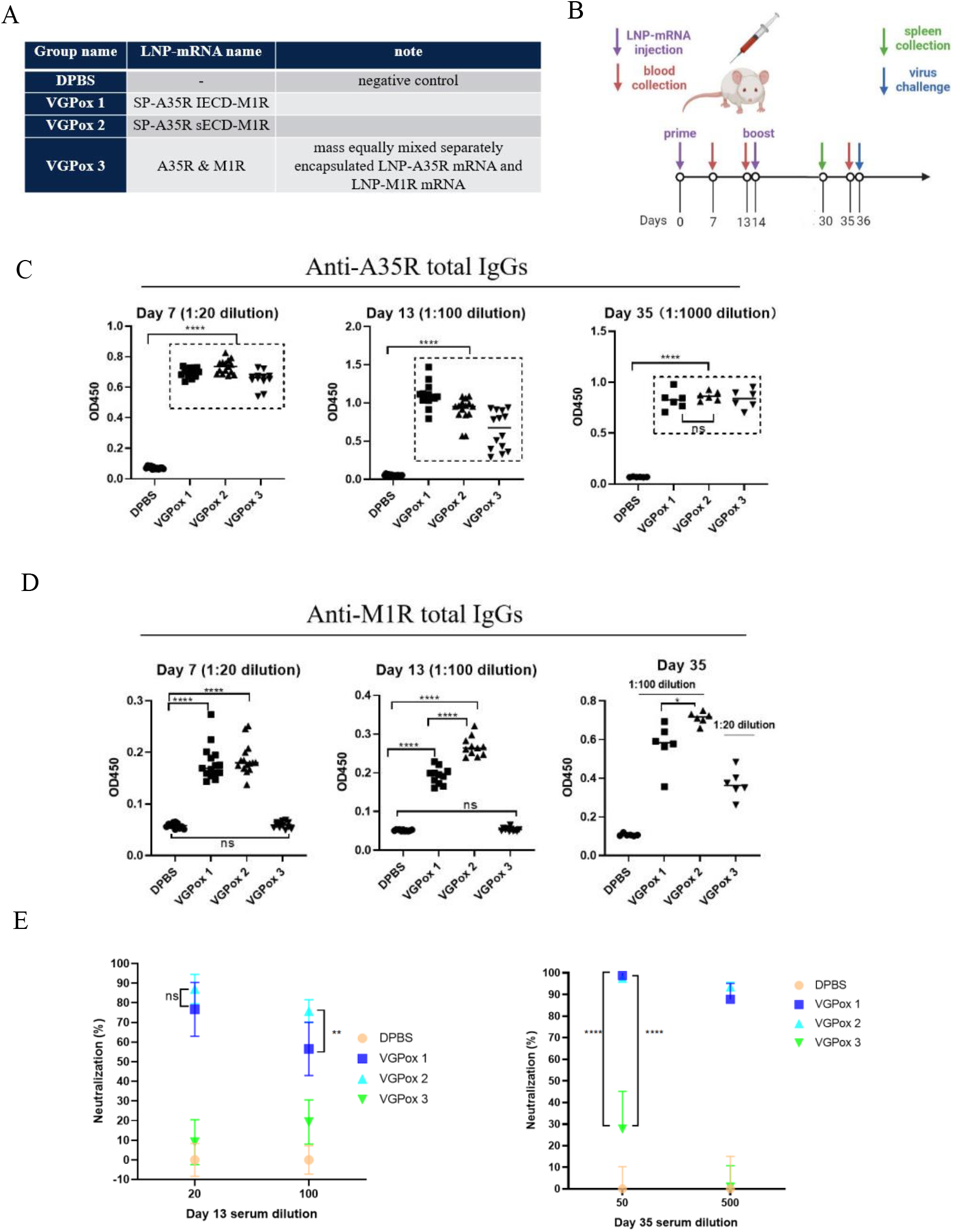
mice vaccination and antibody induction. (A) mice groups. Mice were divided into 4 groups with 15 mice in each group. Mice vaccinated with DPBS were used as negative control. (B) mice vaccination and samples collection. LNP-mRNA was administered intramuscularly at day 0 and 14. Blood was collected at day 7, 13, and 35. One group of mice was sacrificed at day 30 to collect the spleens. Mice were intranasally challenged with a lethal dose (1×10^6^ pfu) of WR vaccinia virus. (C and D) A35R and M1R specific total IgGs, respectively. Serum with different dilutions at different timepoints was used for ELISA against A35R or M1R. (E) Neutralizing antibodies were measured. The sera were diluted and incubated with the virus for PRNT. Serum from DPBS group were regarded as 0% neutralization.

### MRNA vaccines can activate T cell immune response

In order to know whether the designed mRNA vaccines can induce the activation of T cells. Here, we sampled mice spleens at day 30 post vaccination and cells were isolated for Flow cytometry analysis. As shown in Fig. 3A, we used monkeypox A35R homolog vaccinia A33R protein to stimulate CD4 and CD8 T cells by elevation of IFNγ and CD69. It is apparent that both CD4 and CD8 T cells were activated in VGPox 2 and VGPox 3 but not VGPox 1 vaccinated animals. On the other hand, treatment of the spleen cells with a recombinant M1R protein can stimulate CD4 and CD8 T cells in VGPox 1 and VGPox 2 but not VGPox 3 treated animals (Fig. 3B).

**Fig. 3.**
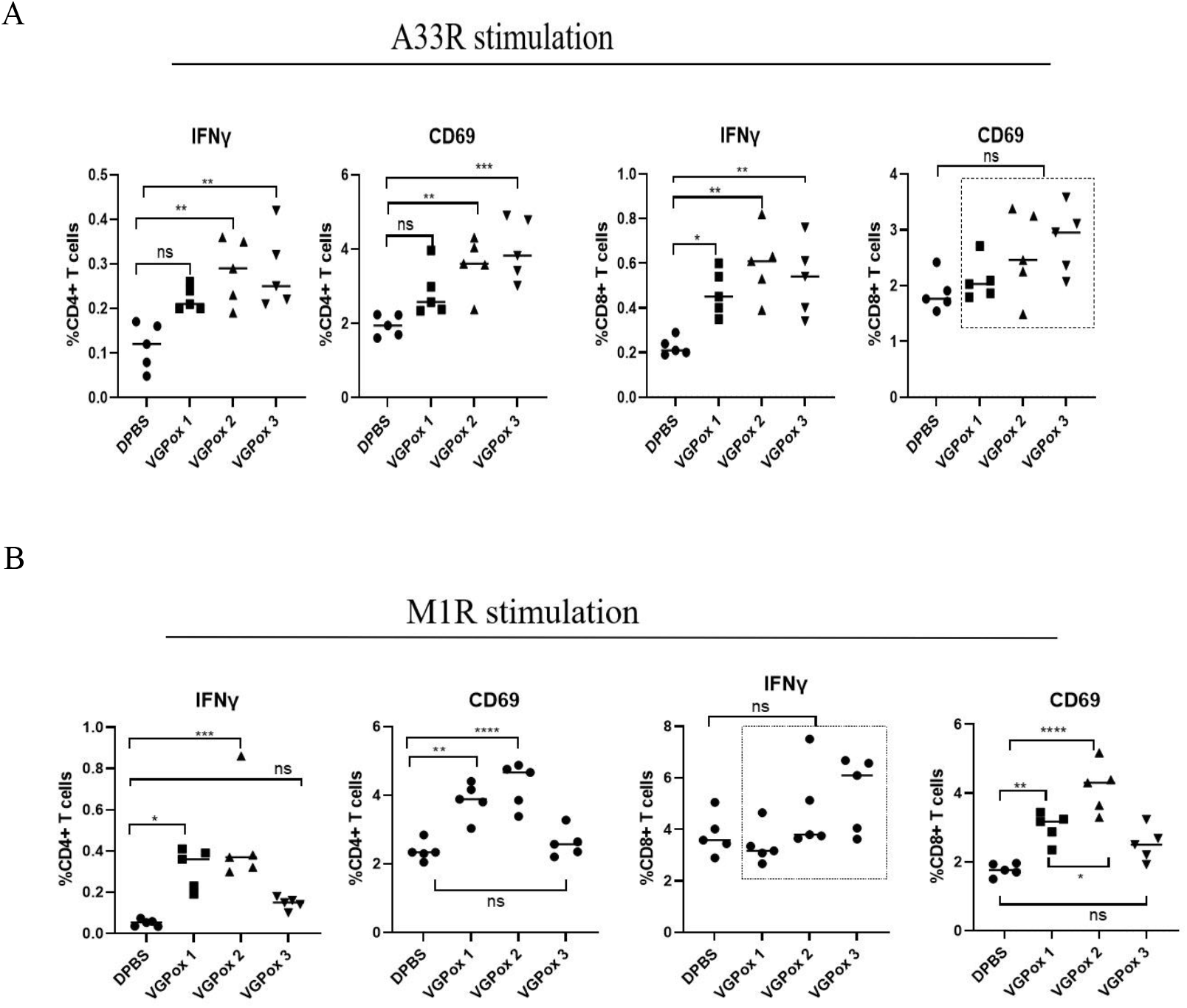
T cell immune response. (A) CD4 and CD8 T cell immune response against A33R. A33R from VACV is the homolog of MPXV A35R. (B) CD4 and CD8 T cell immune response against M1R.

### All 3 mRNA vaccines completely protected mice from a lethal dose virus challenge

To test whether the mRNA vaccines can protect animals from virus infection, vaccinated mice were intranasally challenged with a lethal dose vaccinia virus (VACV-WR). The body weight in DPBS control group started to loss 2 days post infection and nearly lost 15% of the initial weight at day 4. Meanwhile, mice in all mRNA vaccine groups had no body weight loss or other abnormality (Fig. 4A). In consistence to the body weight change, there was a complete virus clearance in lungs of those mice 9 days following nasal inoculation of the virus while the animals in the control group had a high virus load in the lungs. (Fig. 4B).

**Fig. 4.**
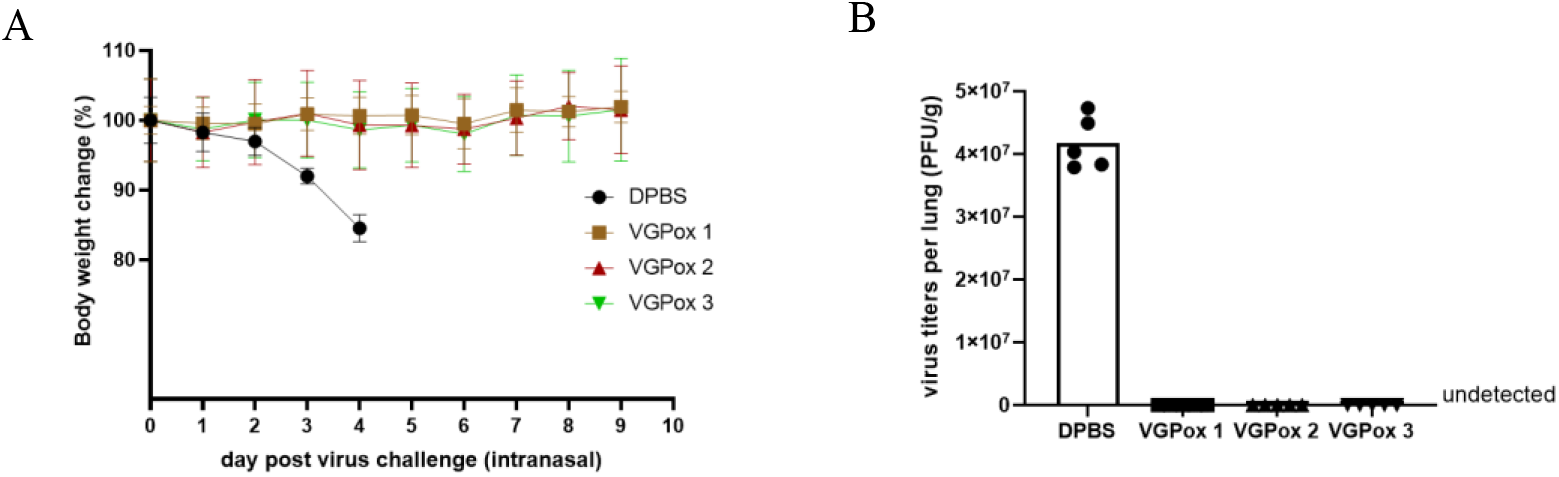
mice protection assay. (A) mice were protected from virus challenge. Body weight of each mouse was examined every day post virus challenge and its changes was calculated compared with initial weight. (B) Viral load in the lungs. The lungs were collected at day 4 when mice lost body weight nearly 15% in DPBS group or at day 9 post virus inoculation in other groups and virus titers were examined by plaque assay.

## Discussion

The current study tested three mRNA vaccines for poxvirus. VGPox 1 and 2 are both single mRNA molecule coding a fusion protein composed of extracellular domain of A35R and a full length M1R, but VGPox 3 contains the mixture of two individual mRNA-LNP complexes coding the A35R and M1R, respectively. The only difference between VGPox 1 and 2 is that the latter further deleted the stalk region of A35R. Our results showed that VGPox 1 and 2 induced much higher antibody levels against M1R than co-expressing the two full-length proteins (VGPox 3) at early time points although the three vaccines produced similar levels of antibodies for A35R. Interestingly, only VGPox 1 and 2 but not VGPox 3 vaccinated sera could neutralize live virus at early time points. Since IMV is the most abundant form of viral particles, it may not be surprised that VGPox 3 vaccine that does not induce high levels of anti-M1R antibodies is not effective for the in vitro neutralization assay.

Comparing between VGPox 1 and 2, VGPox 2 showed higher levels of total IgG against M1R than VGPox 1 and VGPox 3 at all-time points. We noticed that VGPox1 had lower protein expression level than VGPox 2 in 293T cells measured by both anti-A35R and anti-M1R antibodies. This was also confirmed by another anti-A35R antibody (data not shown). It is not clear how much the difference in protein expression level contribute to the different levels of IgGs induced by the two vaccines.

The most striking results of our current study is that the mRNA coding for the fusion forms of A35R and M1R (VGPox 1 and VGPox 2) can effectively induce high levels of both A35R and M1R IgGs and are highly effective in neutralizing live virus infection in cell cultures, but not the mixture of the two individual mRNAs (VGPox 3). VGPox 3 produced M1R specific antibodies much later, and consequently the sera collected at an early time points was not able to neutralize the virus.

Nevertheless, all three mRNA vaccines (VGPox 1-3) are 100% protective in the virus challenge assay. We speculate that this is because that all animals in the current study were challenged with live virus at day 36 when both anti-A35R and anti-M1R neutralizing antibodies were present in all three vaccines. Since neutralizing antibodies against EEV and IMV may be both needed for protection of virus challenge (Gilchuk et al., 2016), it remains to be investigated whether VGPox 1 and VGPox 2 have better protection than VGPox 3 when virus challenge is performed at an earlier time point when the anti-M1R antibodies have not been induced by VGPox 3 vaccine.

In conclusion, our study demonstrated a novel mRNA vaccine expressing a fusion protein composed of monkeypox A35R extracellular domain and M1R can effectively induce both humoral and cellular immunity against the virus, demonstrating a complete protection for the mice with lethal vaccinia virus challenge. Given the high homology of vaccinia and monkeypox, our results suggest that VGPox 1 or 2 can be potential mRNA vaccines against monkeypox.

